# Global distribution and richness of *Armillaria* and related species inferred from public databases and amplicon sequencing datasets

**DOI:** 10.1101/2021.06.29.450419

**Authors:** Rachel A. Koch, Joshua R. Herr

**Author notes:** Correspondence: Joshua R. Herr.

## Abstract

*Armillaria* is a globally distributed fungal genus most notably recognized as economically important plant pathogens that are found predominantly in forest and agronomic systems. The genus *sensu lato* has more recently received attention for its role in woody plant decomposition and in mycorrhizal symbiosis with specific plants. Previous phylogenetic analyses suggest that around 50 species are recognized globally. Despite this previous work, no studies have analyzed the global species richness and distribution of the genus using data derived from fungal community sequencing datasets or barcoding initiatives. To assess the global diversity and species richness of *Armillaria*, we mined publicly available sequencing datasets derived from numerous primer regions for the ribosomal operon, as well ITS sequences deposited on Genbank, and clustered them akin to metabarcoding studies. Our estimates reveal that species richness ranges from 50 to 60 species, depending on whether the ITS1 or ITS2 marker is used. Eastern Asia represents the biogeographic region with the highest species richness. We also assess the overlap of species across geographic regions and propose some hypotheses regarding the drivers of variability in species diversity and richness between different biogeographic regions.

## INTRODUCTION

While the importance that fungi play as mutualists, decomposers, and pathogens is undisputed, researchers are just beginning to disentangle the processes that shape their global species richness and distribution (Tedersoo et al., 2014). One such globally important group is the armillarioid clade—a lineage composed of the mushroom-forming genera *Armillaria* and *Desarmillaria*, along with the sequestrate genus *Guyanagaster. Armillaria* species, representative of the annulate mushroom-forming species within the armillarioid clade, are globally distributed with an estimated 40 to 50 species (Volk & Burdsall, 1995; Baumgartner et al., 2011). In contrast, *Desarmillaria* – which was recently separated from *Armillaria* on the basis of morphological and molecular characters (Koch et al., 2017; Heinzelmann et al., 2019) – is representative of the exannulate mushroom-forming species within the clade. Within *Desarmillaria*, there are three described species, including *Desarmillaria ectypa*, which is restricted to northern European bogs (Zolciak et al., 1997), *D. tabescens*, which is likely found widely throughout Europe and Asia (Guillaumin et al., 1993), and *D. caespitosa*, which is found throughout North America (Antonín et al., 2021). While *Guyanagaster* – representative of the sequestrate species – is a member of the armillarioid clade (Henkel et al., 2010; Koch et al., 2017), herein the discussion of the armillarioid clade will only refer to species of *Armillaria* and *Desarmillaria*, and when necessary, *Guyanagaster* will be specifically referenced.

Armillarioid species are most often recognized as ecologically and economically important plant pathogens, identified as the causal agents of Armillaria Root Rot. As facultative necrotrophs, these species possess both parasitic and saprotrophic phases. During their parasitic phase, species may attack a broad range of woody dicotyledonous hosts in both natural and agronomic systems (reviewed in Baumgartner et al., 2011). However, during their saprotrophic phase, armillarioid species play an important role in carbon cycling and nutrient exchange as they contain a suite of genes that encode for enzymes that have the ability to degrade the entire spectrum of the major components of plant cell walls (Kile et al., 1991; Sipos et al., 2017). Currently, all armillarioid taxa are recognized as having the ability to be a pathogen during their life cycle – even the putatively saprotrophic *A. cepistipes* has been recently reported to be pathogenic on *Populus trichocarpa* (Alveshere et al., 2021). One largely diagnostic characteristic of many armillarioid species is the production of rhizomorphs – discrete, long-lasting filamentous structures that forage for new resources away from the primary resource base (Garraway et al., 1991). Rhizomorphs can grow long distances to infect plants, resulting in genets with exceptionally large sizes, some of which have been recognized as the largest and oldest organisms on earth (Smith et al., 1992).

In addition to their importance as plant pathogens and biomass decomposers, armillarioid species are ecologically important as they have the capacity to engage in several symbiotic associations with various plants, other fungi, and insects. For example, the growth of sclerotia of the polyporoid fungus, *Polyporus umbellatus*, is dependent on nutrition from symbiotic *Armillaria* species (Guo & Xu, 1992). *Armillaria* species in east Asia engage in a form of mycorrhizal relationship known as mycoheterotrophy with species of achlorophyllus orchids in the genera *Galeola* and *Gastrodia* (Kikuchi & Yamaji, 2010; Guo et al., 2016). Arguably, *Armillaria* sporocarps may be one the most documented mushroom target of other pathogenic fungi as they are specifically parasitized by the mushroom-forming species *Entoloma abortivum* (Lindner Czederpiltz et al., 2001; Koch & Herr, 2021) and putatively by species in the Ascomycete genus *Wynnea* (Xu et al., 2019). Lastly, the sequestrate genus *Guyanagaster* is the earliest diverging lineage within the armillarioid clade (Koch et al., 2017). The sporocarps of *G. necrorhizus* house nitrogen-fixing bacterial species, which likely facilitates a dispersal mutualism with wood-feeding termites (Koch et al., 2018; Koch et al., 2021).

Traditionally, species within the armillarioid clade have been delimited by morphological characteristics and, more recently, by mating compatibility studies (Bérubé & Dessureault, 1989; Cha et al., 1994; Volk & Burdsall, 1995; Qin et al., 2007). However, given the difficulty in identifying species based solely on these features, there has been significant emphasis on understanding species boundaries within the armillarioid clade using phylogenetic data (Guo et al., 2016; Klopfenstein et al., 2017; Koch et al., 2017; Coetzee et al., 2018). Initially, studies of the armillarioid clade, like most fungal diversity assessments, relied heavily on data from the nuclear ribosomal operon – which bridges the small subunit (18S), internal transcribed spacer region 1 (ITS1), 5.8S subunit, internal transcribed spacer region 2 (ITS2), the intergenic spacer region (IGS), and the large subunit (28S) (Chillali et al., 1998). However, the *translation elongation factor 1* (*tef1*) is the marker region that has emerged in more recent research on armillarioid fungi, in that it provides adequate resolution to infer species boundaries and lineage relationships within a global context (Maphosa et al., 2006; Guo et al., 2016; Klopfenstein et al., 2017; Coetzee et al., 2018). While the research community interested in armillarioid fungi has recently focused on the use of the *tef1* marker, this locus has not been utilized in any environmental sequencing studies with which to measure global diversity and richness. In order to evaluate both diversity and richness on a global scale, it is ideal to have a suite of characters – including both morphological and molecular data – from many different sampling locations. While the mycological research community is slowly moving toward the collection of molecular data from multiple markers on a global scale, research has so far focused solely on characterizing fungal species richness using the ITS1 and ITS2 regions that have consensus from the fungal research community (Schoch et al., 2011; Herr et al., 2013; Hibbett et al., 2015).

Given the importance of armillarioid fungi, we wanted to understand the diversity and richness of this group on a global scale. To do this, we assess – from the perspective of the widely adopted in environmental barcoding markers, ITS1 and ITS2 – the overall global distribution and species richness within this clade, much like has been done previously for soil fungi on a large scale (Tedersoo et al., 2014). Here, we analyzed the totality of publicly available ITS sequences submitted to NCBI-GenBank and fungal community sequencing data using metabarcoding methodologies submitted to NCBI-SRA to gain a more complete understanding of the species richness and the global distribution of the armillarioid clade. We recognize that it is important to understand how this locus operates for this lineage given the immense number of studies utilizing it, and we hypothesize that the ITS1 and ITS2 regions exhibit concordance. We also discuss the utility of public data to inform species clusters, in this case operational taxonomic units (OTUs), in armillarioid fungi and present caveats to its use. Additionally, we speculate on factors, such as symbiotic interactions of armillarioid species, that affect the diversity and geographic distributions of this lineage.

## MATERIALS AND METHODS

### Sequences curated from public database vouchers

To create a dataset of all publicly available armillarioid ITS sequences, we queried NCBI-GenBank (https://www.ncbi.nlm.nih.gov) for “*Armillaria*” with “ITS” (or “*Desarmillaria*” and “*Guyanagaster*” with “ITS”) and downloaded all of the nucleotide sequences matching these criteria. Taxonomy Browser (NCBI) (Federhen, 2012), which uses BLAST+ (Camacho et al., 2009), was then used to verify the correct taxonomic names – those entered by the researchers who deposited the sequences to NCBI – for each of the public sequences. We did this to remove sequences that were deposited to NCBI under an incorrect genus-level designation (Bruns et al., 2008). The feature for geographic region (such as “country”) was also extracted for each sequence and any sequences without this corresponding metadatum were excluded. Additionally, if sequences were associated with another organism in a symbiotic relationship, this was also noted. Finally, all sequences with the identifier “*Armillaria*” from the UNITE v8.3 database (Nilsson et al., 2018) were also included in the dataset irrespective if they had corresponding location metadata. For these sequences, we then extracted the ITS1 and ITS2 regions using ITSx (Bengtsson-Palme et al., 2013).

### Sequences curated from public amplicon studies

We next mined the Sequence Read Archive (SRA) database for public datasets of fungal amplicon sequencing data of the ITS1 and ITS2 region. We identified 454 pyrosequencing and various Illumina platform sequencing datasets with the query (ITS2, ITS1 AND “biomol dna”[Properties] AND “platform ls454”[Properties] AND “platform Illumina”[Properties]) to the SRA database (Leinonen et al., 2010) which provided us with 183,838 sample files representing 1,253 public deposits to the SRA. We then manually curated a list of datasets based on the availability of metadata from the NCBI SRA Run Selector, such as geographic location, as we did previously for the NCBI sequences. Once datasets were downloaded in the “.sra” format using the ‘prefetch’ command, we converted the datasets to “.fastq” format using the ‘fasterq-dump’ command with the “--split_files” option. Both tools are available from NCBI through the sra-tools scripts (https://github.com/ncbi/sra-tools). Sequences from SRA deposited datasets were clustered against the NCBI list for *Armillaria* (generated in the first stage) using VSEARCH version 2.17.0 (Rognes et al., 2016) at 90% sequence similarity using the ‘--usearch_global’ command. To avoid off-target clustering and to double check the VSEARCH clustering quality, we next identified the output from the VSEARCH clustering using DECIPHER version 3.13 IDTAXA algorithm ‘Classify Organisms’ (Wright, 2016) and the UNITE2020 DECIPHER database (http://www2.decipher.codes/Classification/TrainingSets/UNITE_v2020_February2020.RData) selecting only those sequences being taxonomically identified as *Armillaria* and *Desarmillaria*. Out of the 1,253 public studies deposited to the SRA that we queried, only 56 studies, restricted to forest soil and plant matter, contained sequences with 90% homologous base pairs to *Armillaria* ITS sequences. The remaining 1,197 studies, which mainly consisted of ecosystems that did not contain woody plant hosts (i.e. built-environment, human, marine, etc.), did not contain any sequences with homology to existing *Armillaria* sequences.

### Curation of total sequence datasets and geographic metadata

Geographic location metadata were also extracted from studies containing *Armillaria* sequences, which represented *Armillaria* voucher specimens deposited in NCBI Genbank and from the SRA database metadata files. Sequences of the ITS1 region from the Genbank nucleotide database and the SRA were combined, and the same was done as a separate dataset for the ITS2 region. Any sequences less than 150 base pairs in length were removed. To estimate species from these sequences, we then clustered the ITS1 and ITS2 regions (combining the voucher specimen data and SRA sequences) at 98% using USEARCH 11 (Edgar, 2010). We chose 98% nucleotide identity threshold because this was the identity threshold utilized by Tedersoo et al. (2014). The biogeographic regions we refer to largely follow those utilized by Tedersoo et al. (2014). Because some of the armillarioid sequences associated with *Wynnea* species did not pass our quality filters and we were still interested in utilizing them, we independently placed each of them into our dataset using BLAST+, as above, to place them in the appropriate species cluster.

### Phylogenetic analysis

To visualize how the armillarioid species clusters were related, we utilized a phylogenetic approach. One exemplar sequence from each species cluster was chosen to include in the multiple sequence alignment. We utilized the full-length sequence, when possible, and not just the hypervariable regions. Species clusters for the ITS1 region and ITS2 region were analyzed separately. This was because in no study where we identified armillarioid taxa did researchers employ both ITS1 and ITS2 markers from a single sampling location such that we could absolutely ensure that both markers came from the same genetic individual or genet. As ITS1 and ITS2 sequences could not be ascribed to a single genet or individual, we did not attempt to statistically assess the phylogenetic tree correlation of the ITS1 and ITS2 sequences – this would be better suited for pangenomic studies of the armillarioid fungi using multiple genes. The public sequences we identified as having homology to our armillarioid nexus file were aligned using MAFFT version 7 (Katoh & Standley 2013) with refinements to the alignment performed manually. RAxML-NG (Kozlov et al., 2019) was used to reconstruct this phylogeny. The two known species of *Guyanagaster*, sister to *Armillaria* and *Desarmillaria* species (Koch et al., 2017), were used to root this phylogeny. We defined well supported branches as having 70% or greater bootstrap support. The location metadata associated with each species cluster was mapped onto each phylogenetic tree using the ‘ggtree’ package (Yu et al., 2018) within the R 4.1.0 environment (R Core Team 2013).

## RESULTS

A total of 1,153 *Armillaria* ITS1 region sequences (1,119 from the nucleotide database and 34 from the SRA database) and 1,567 *Armillaria* ITS2 region sequences (1,246 from the nucleotide database and 321 from the SRA database) were sorted into species clusters. Public data sets which contributed to the high-throughput ITS1 and ITS2 region sequences in this study consisted of six different primer pair combinations: fITS7/ITS4, gITS7/ITS4, ITS1F/ITS2, ITS1F/ITS4, ITS3/ITS4, and ITS5/ITS2. Sequences for both ITS marker regions were identified from samples collected from all continents except Antarctica. At a 98% nucleotide identity, 51 armillaroid species clusters were recovered for the ITS1 region and 61 for ITS2 region. The geographic distribution of these species clusters are shown in Fig. 1 and the symbiotic associations of these species clusters is displayed in Table 1. Of the ITS1 region species clusters, sequences from the SRA database clustered into five species clusters (species clusters 5, 6, 9, 15 and 23), all of which also contained sequences queried from NCBI-Genbank. Of the ITS2 species clusters, sequences derived from the SRA database were clustered into 14 species at 98% similarity (species clusters 3, 9, 10, 13, 17, 18, 19, 23, 24, 26, 30, 35, 43 and 51); one of these clusters, species cluster 17, was solely composed of sequences from the SRA database and did not cluster with any sequences from the NCBI nucleotide database.

**Fig. 1.**
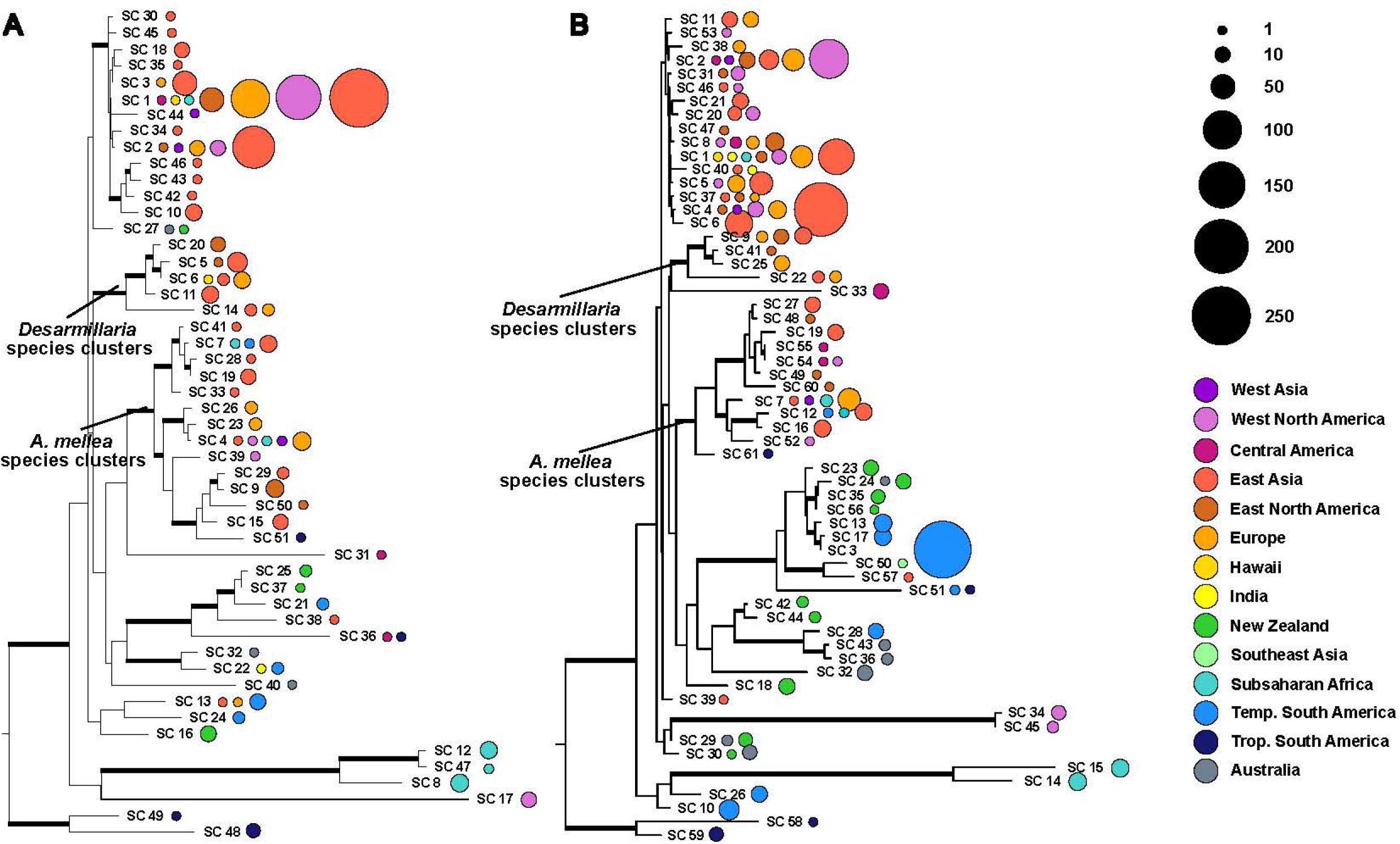
Phylogenetic trees illustrating the number of sequences that correspond to each species cluster. Maximum likelihood phylogeny constructed from the full length ITS region using one exemplar sequence per species cluster. Two species of *Guyanagaster*, sister to all other armillarioid species, were used to root this phylogeny. Well-supported branches (greater than 70% bootstrap support) are thickened. The color of the corresponding circles corresponds to the biogeographic region from which the voucher material producing the sequences was collected. The size of the circles is scaled by the number of sequences that were grouped into a particular species cluster from a particular region. (A) Phylogeny of species clusters as determined from analysis of ITS1 and (B) phylogeny of species clusters as determined from analysis of ITS2.

**Table 1.**
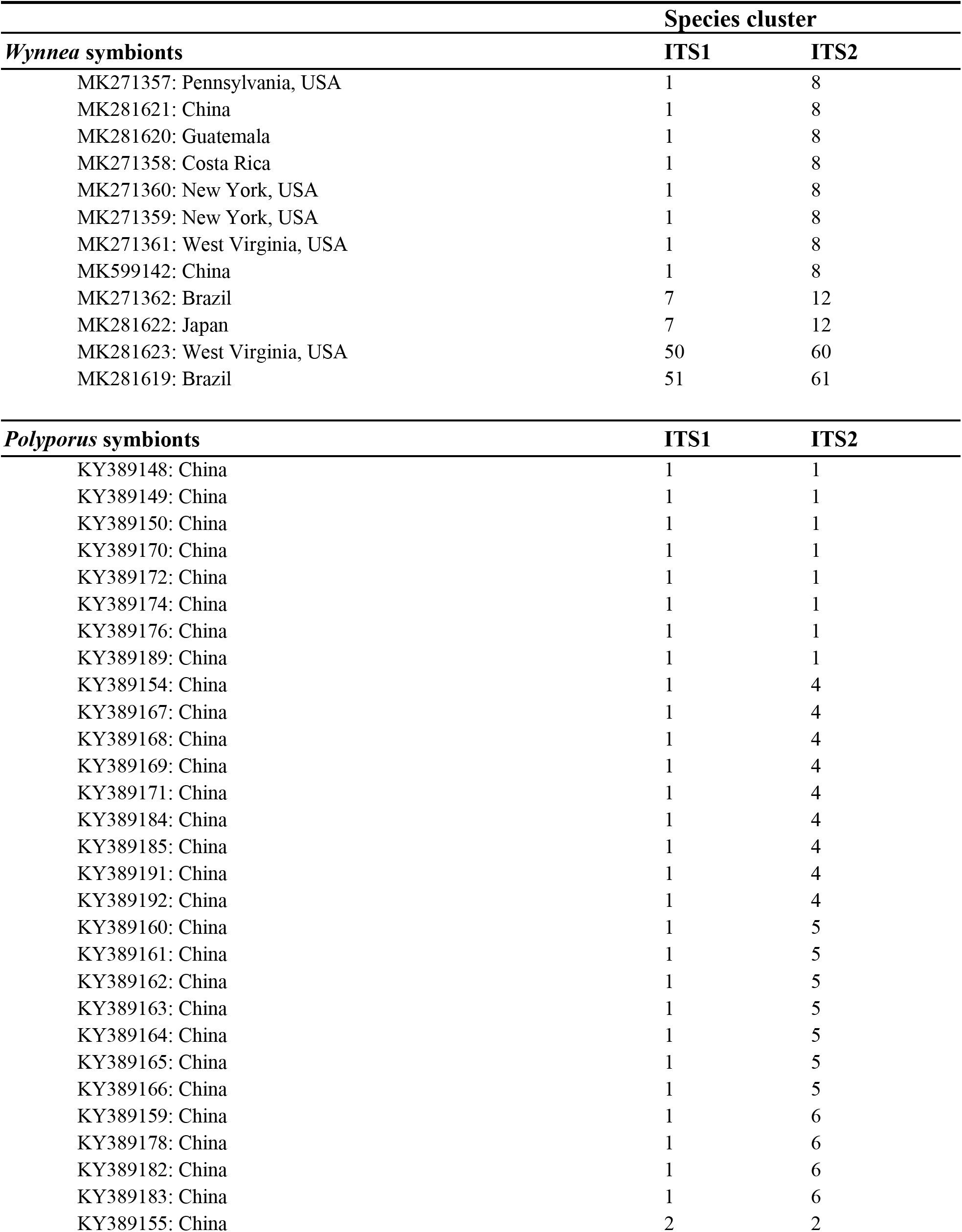
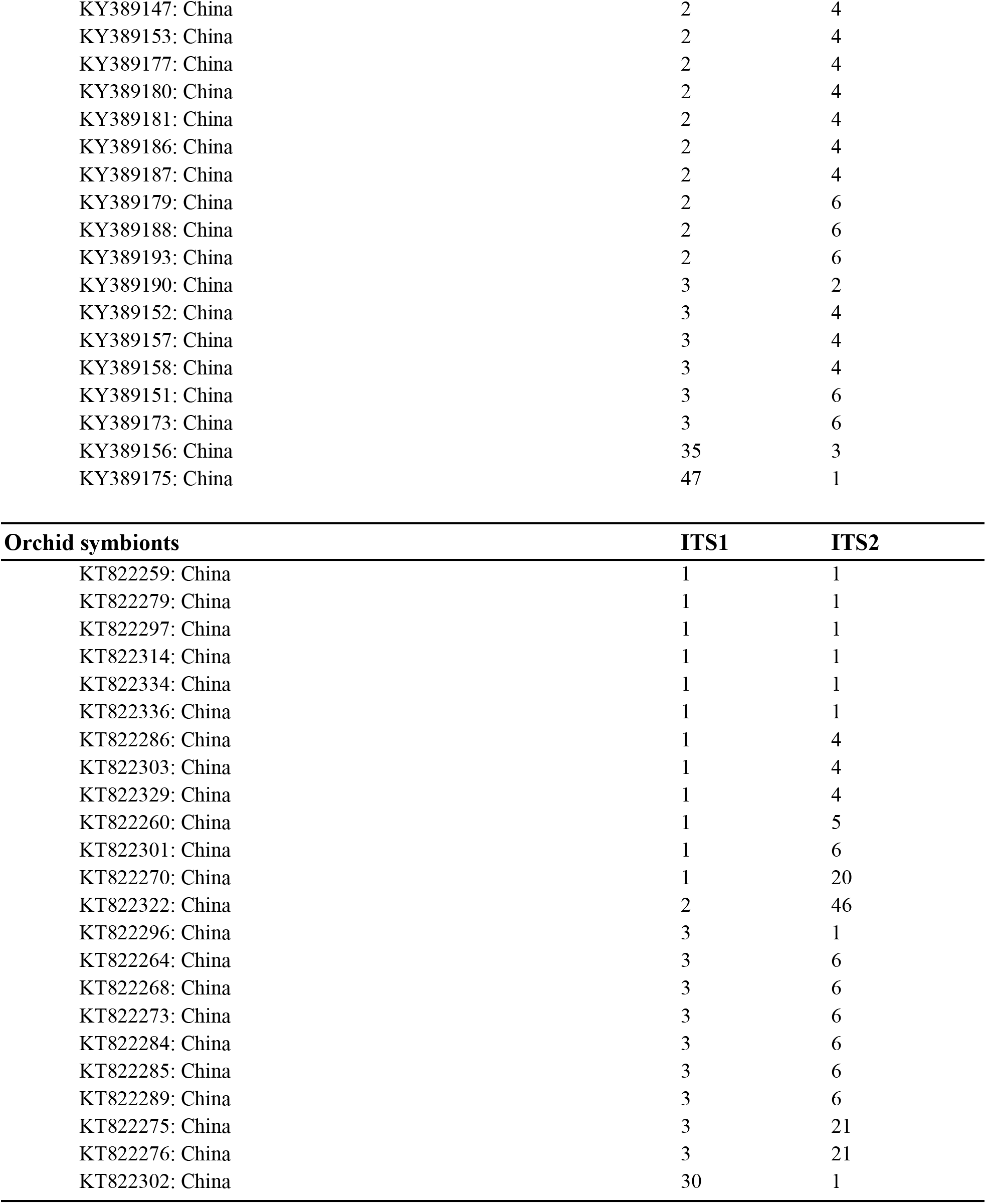
Armillarioid sequences (and corresponding GenBank accessions and location metadata) from tissue engaging in symbiotic associations.

The most species rich biogeographic region was East Asia, with 23 and 19 species clusters at ITS1 and ITS2, respectively (Fig. 1 and 2). With the ITS1 region data, East Asia also contained the most species clusters that are found in no other biogeographic region (Fig. 2). The other species rich biogeographic regions included western and eastern North America and Europe, with five, six, and nine armillarioid species clusters at the ITS1 region and 13, 12, and 12 armillarioid species clusters at the ITS2 region, respectively. These geographic regions appear to be the best sampled as a majority of the total barcoded armillarioid sporocarps, rhizomorphs, and mycelial tissue are from these regions (Fig. 3). In the southern hemisphere, species richness is lower when compared to the northern hemisphere, with temperate South America, Africa, Australia, and New Zealand having five, six, three, and four species clusters at the ITS1 region and eight, five, six, and nine species clusters at the ITS2 region. Largely tropical geographic areas, including Hawaii, India, tropical South America, and Southeast Asia have the lowest species richness with one to two mushroom-forming species in each location. Tropical South America also has two species of the sequestrate genus, *Guyanagaster*.

**Fig. 2.**
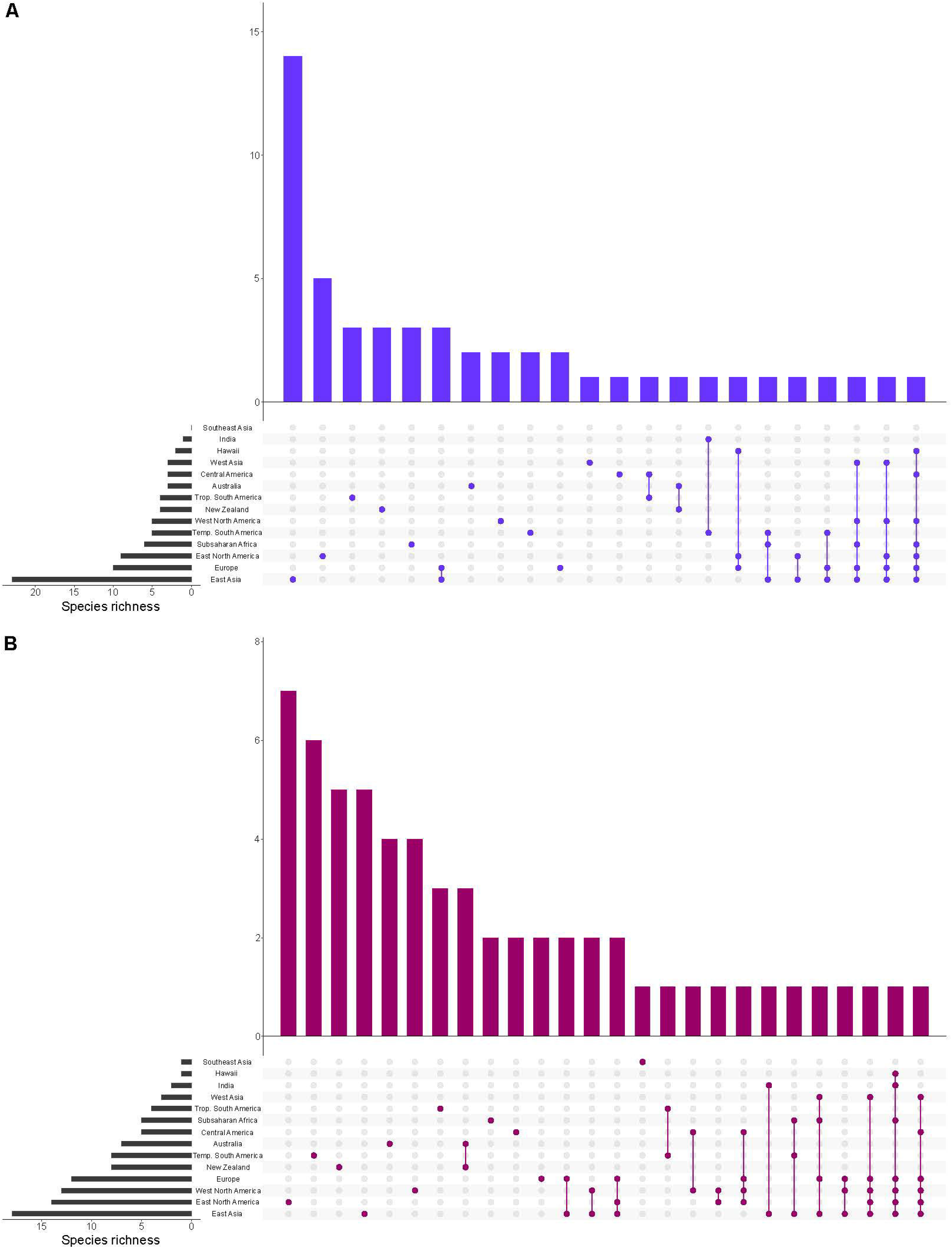
Species richness in each of the biogeographic regions. Bar graphs showing the total richness of our species clusters in each biogeographic region and the combination of biogeographic regions. (A) ITS1 and (B) ITS2. The species richness for each vertical bar corresponds to the number of species that are solely found in the biogeographic regions corresponding to the colored circle(s) below. The species richness for the horizontal insets correspond to the total species richness within each biogeographic region.

**Fig. 3.**
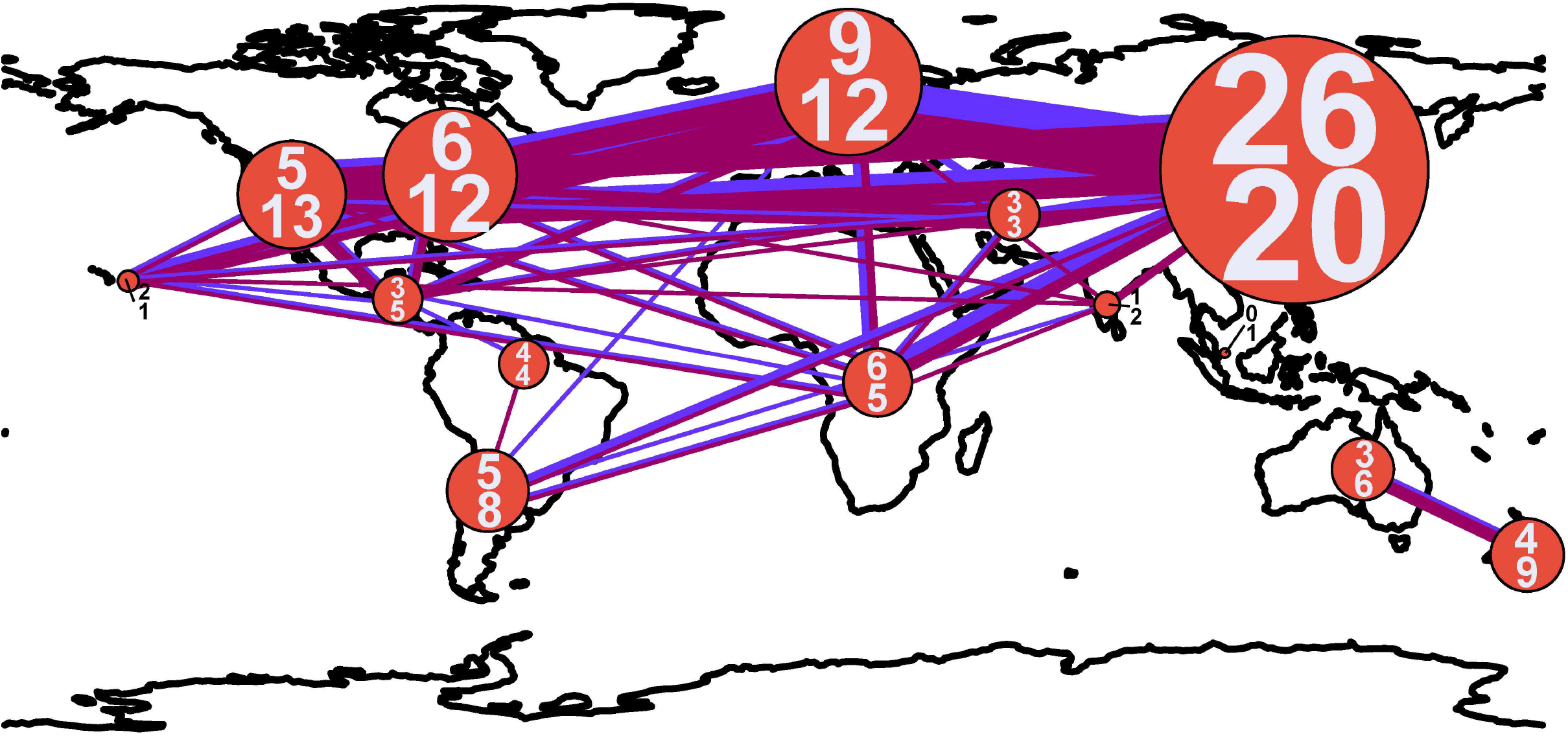
Connectedness of biogeographic regions by shared armillarioid species clusters. The width of the lines and diameter of circles are proportional to the number of connections and number of species, respectively. The blue line corresponds to ITS1 and the magenta line to ITS2. The numbers within the circles indicate the number of armillarioid species clusters detected in each region, with the top number referring to the number of species clusters at ITS1 and the bottom number at ITS2.

Shared species clusters between biogeographic regions were much more pervasive in the northern hemisphere compared to the southern hemisphere. Between locations, the most shared species clusters include a link between East Asia and Europe, as well as a link between Australia and New Zealand. Evidence for shared species clusters throughout the southern hemisphere is limited to only Australia and New Zealand (Figure 2), but according to our analysis, these regions do not share any species clusters with other geographic location (Fig. 1). At the ITS1 region, four species clusters (species clusters 1, 4, 13, and 22), and at the ITS2 region, three species (species clusters 1, 7, and 12), were shared between locations in the northern and southern hemisphere. Two of the species clusters derived from the ITS1 region and all of the species clusters from the ITS2 region included sequences from specimens or environmental samples in the northern hemisphere and Subsaharan Africa.

Species clusters (or species complexes) for the armillarioid lineage were determined from the combined dataset of sequences derived from community sequencing efforts and accessions deposited into NCBI Genbank. We identified two known sequestrate armillarioid species— species of *Guyanagaster*—from the nucleotide database. Of the *Desarmillaria* species, there was one species cluster detected at both the ITS1 (species cluster 14) and ITS2 (species cluster 22) regions, from Europe and East Asia, which likely corresponds to *D. ectypa*. Four and three species clusters that correspond to other *Desarmillaria* species at the ITS1 (species clusters 5, 6, 11 20) and ITS2 (species clusters 9, 25 and 41) regions, respectively, were also detected. The *Desarmillaria* species clusters form a well-supported monophyletic lineage in analyses of both ITS1 and ITS2 marker regions. All of the other species clusters, 45 at the ITS1 region and 56 at the ITS2 region, likely correspond to *Armillaria sensu stricto*. There were 12 species clusters at the ITS1 (species clusters 4, 7, 9, 19, 23, 26, 28, 29, 33, 39 and 41) region and ten at the ITS2 (species clusters 7, 12, 16, 19, 27, 48, 49, 52, 54 and 55) region that included sequences annotated as *A. mellea*. These *Armillaria mellea* species clusters, along with two *Wynnea* symbionts (from both an ITS1 and ITS2 marker region perspective) that likely represent as-yet undescribed species, form a well-supported monophyletic lineage in both analyses (Fig. 3), indicating that this is a genuine species complex as illustrated in the phylogenetic trees in Fig. 3. Species clusters for both the ITS1 and ITS2 regions are available in Tables S1 and S2.

## DISCUSSION

Previous studies within the armillarioid clade have used morphology (Bérubé & Dessureault, 1988; Watling et al., 1991; Volk & Burdsall, 1995), hyphal compatibility tests (Anderson & Ullrich, 1979; Banik & Burdsall, 1998) and phylogenetic assessment (Maphosa et al., 2006; Guo et al., 2016; Coetzee et al., 2018) to draw conclusions of diversity and global richness. In this study, we utilized a complementary methodology — data derived from both the ITS1 and ITS2 marker regions taken from barcoding and community sequencing analysis studies — to investigate global species richness and distribution of species. As the universal barcode for fungi (Schoch et al., 2013), the ITS regions are the most widely available in terms of number of species that are represented in public databases (Nilsson et al., 2018), and the region is the predominant marker for fungal molecular ecology studies (Herr et al., 2013, Hibbett et al., 2016). Additionally, there has been an increasing awareness of the importance of barcoding macroscopic fungal collections, most notably those stored in fungal herbaria, using ITS primers to aid in identification and create more complete databases (Truong et al., 2017; Haelewaters et al., 2018; Vu et al., 2019). Therefore, the strength of this approach is that it allowed us to utilize the increasing numbers of publicly available datasets (notably using the ITS1 and ITS2 marker regions) from fungal ribosomal operon barcoding and metabarcoding efforts that have previously gone unused in *Armillaria* phylogenetic studies.

The data we analyzed for the ITS1 region suggests there are 51 species clusters in the armillarioid clade, while the ITS2 region data indicates there are 61 species clusters. These numbers are largely consistent with what has been previously determined through multigene phylogenetic efforts focused largely on fungal sporocarp tissue samples (Guo et al., 2016; Coetzee et al., 2018). Given our filtering scheme for short and spuriously aligned sequences, and the fact that we did not include any singletons – recognized here as species clusters here that only contained a single sequence – our results have likely underestimated the species richness of this clade. Additionally, as the number of environmental sequencing studies increase, particularly in under-sampled regions across the globe, we will likely uncover an even greater species richness. In additional to increased geographic sampling, future studies may also find – when using a greater number or a more phylogenetically informative marker region – that overall species richness of the armillarioid clade is greater than we report here.

There are known biases with ITS as a marker within the armillarioid clade, including that it may be too conserved to discriminate some species (Chillali et al., 1998; Coetzee et al., 2001). Our findings corroborate this, particularly when it comes to discriminating between *A. gallica*, *A. sinapina*, *A. cepistipes*, *A. calvescens* and *A. ostoyae*. (Figure 3). Both the ITS1 and ITS2 marker regions failed to resolve these taxa with any certainty. However, the other possibility for this observation is not because of poor resolution from the markers, but because of a misapplication of names deposited into GenBank (Bruns et al., 2008). In other instances, the ITS regions are also too variable and may lead to an overestimation of individuals in other species complexes (Dunne et al., 2002). This may certainly be the case with *A. mellea*, which could represent up to 14 species based on analysis of the ITS1 region (Figure 3). While our results suggest that ITS1 and ITS2 may provide estimates of species richness that are consistent with morphological, reproductive behavior, and multigene phylogenetic assessment, these markers are unlikely to provide an accurate reflection of precise delimitation in the armillarioid fungi. Species delimitation is best employed with large datasets with multiple different characters and suites of genes – we have found that this is most certainly the case with the armillarioid fungi. This study, which is based on two public marker regions using seven primer pairs, underscores the need for additional ecological markers for fungi and for the development of sequence analysis from fungal metagenomic and metatranscriptomic studies (Kuske et al., 2015; Hibbett et al., 2016). However, because the aim of this work is to understand diversity and distribution patterns, and not to resolve taxonomy issues within this clade, at present we are limited by species clusters identified by our marker regions and taxonomic identification derived from previous phylogenetic analyses and the expertise and reliability of data submitters on Genbank. We argue that further taxonomic work is accomplished via a combination of morphological, ecological, and phylogenomic data.

Reconciling species clusters – which we identify as OTUs – and separating species boundaries is challenging problem in microbial ecology, taxonomy, and phylogenetics (Mysara et al., 2017). The species clusters we generated here, analogous to the reference sequences in the UNITE database for fungal ITS, were generated by clustering homologous sequences at 98.0% similarity (Nilsson et al., 2018). Like the UNITE database, as well as other databases for the ribosomal operon, our clustering is a close approximation to species but arbitrary in designation. We rely here on manual curation to choose a species designation for a particular cluster or OTU. We have roughly designated species in Figures 1 and 2, but these should only be considered accurate when multiple factors, such as multi-locus or genomic sequence data, as well as morphological and mating studies are incorporated (Hibbett et al., 2016).

### Patterns of distribution and species richness

From the perspective of the ITS1 marker region, all armillarioid sequences derived from our curated SRA-derived sequences clustered with existing sequences from the nucleotide database based on voucher collected material. The ITS2 marker region is utilized more frequently in fungal community sequencing efforts due to its level of variability (Hibbett et al., 2016), so it is not surprising that we detected more armillarioid species clusters when using this marker region. The sequences from our curated amplicon sequencing dataset that are not represented by voucher-derived sequence reads represent a single armillarioid species cluster from temperate South America, a geographic area warranting increased surveys for sporocarp diversity. Because several *Armillaria* species from that region do not have sequenced type or voucher specimens, it remains unknown if our armillarioid species cluster represents an undescribed species or represents a species that has been described in the literature but has not been previously sequenced. Another interesting example, and one that elucidates the importance of barcoding studies, is the species cluster 5 from the ITS1 region which is composed of one sequence from a voucher specimen (Vu et al., 2019) and all the other sequences being from an environmental study with no associated voucher material. This further illuminates the importance of continued efforts to barcode, by using multiple marker regions or genomic sequencing, well documented voucher material.

As an important pathogen in forest and agronomic systems there has been an emphasis on providing data to advance species delimitation of *Armillaria.* Additionally, the relatively characteristic signs of *Armillaria* presence in the environment, including white mycelial fans under tree bark and black shoestring-like rhizomorphs (Baumgartner et al., 2011) are easy to identify, collect, culture, and sequence species of *Armillaria*, which certainly aids in our understanding of the distribution of these species even in the absence of sporocarps. Lastly, the entirety of the armillarioid clade has relatively few species when compared to other genera found in the Agaricales (Liimatainen et al., 2014; Oliveira et al., 2020). Of course, the available community sequencing studies from which we mined do not comprehensively represent all natural communities in which armillarioid species occur, so there are undoubtedly more species that are left to be found as sampling increases. However, these results also suggest that *Armillaria* species are not contributing substantially to the pool of species known based purely on sequence data (Hawksworth and Lücking, 2017).

Our analyses here largely agree with previously phylogenetic assessments on diversity of this clade and which geographic regions harbor the greatest species richness in the armillarioid clade. This study shows that armillarioid species richness is concentrated in the northern hemisphere, specifically in East Asia. While this finding can certainly be attributed in part to an outsized collecting effort of this group throughout this geographic area (Fig. 3), it may also be reflective of the life history of this lineage. Eurasia is hypothesized to be the ancestral range of *Armillaria* (Koch et al., 2017) and these results are consistent with this region also being the center of species diversity. East Asia, and China in particular, was previously known to have a high species richness of *Armillaria* species (Qin et al., 2007; Zhao et al., 2008; Guo et al., 2016). Furthermore, phylogenetic analyses have provided significant evidence of genetic divergence between *Armillaria* specimens collected from China that were thought to be conspecific throughout the northern hemisphere, suggesting that endemicity may be prevalent in the northern hemisphere in this clade (Guo et al., 2016) and which our analyses also agree with (Fig. 2).

Another feature of northern hemisphere diversity of the armillarioid clade is the high degree of connectedness between the three primary geographic regions (Fig. 1). Certain *Armillaria* species clusters in the northern hemisphere appear to be broadly distributed throughout the entire region. There are three and five *Armillaria* species clusters, respectively, that are present in North America, Europe and East Asia when considering the ITS1 and ITS2 markers. These species clusters are largely in circumscription with existing designations of the *A. gallica*, *A. calvescens*, *A. sinapina*, and *A. cepstipes* morphological and molecular species complexes. This finding supports the notion that some armillarioid species are distributed throughout the entire northern hemisphere (Coetzee et al., 2015; Coetzee et al., 2018). However, within the northern hemisphere, there are also many species that appear to be restricted to just a certain biogeographic region, suggesting that a narrower interpretation of specific species is more appropriate (Figure 2). For example, the name *A. mellea* have been applied to specimens collected throughout the northern hemisphere. However, our results suggest that what is typically identified as *A. mellea* is in fact a broad species complex (Figure 3). This confirms a previous study which detected significant phylogenetic structure in specimens matching the description of *A. mellea* (Coetzee et al., 2000). While the same has been suspected for *D. tabescens*, inconclusive results from mating studies (Darmono et al., 1992; Guillaumin et al., 1993; Ota et al., 1998; Qin et al., 2007) and phylogenetic analyses (Tsykun et al., 2013; Coetzee et al., 2015), have clouded efforts to disentangle this complex. However, a recent study has separated the *D. tabescens* species complex into two separate taxa based on a five gene phylogeny of voucher collections previously identified as *D. tabescens* (Antonín et al 2021). Our results suggest that there may be even more species in this complex.

Our geographic assessment of diversity is in agreement with numerous other studies that found a distinct suite of *Armillaria* species throughout the southern hemisphere (Coetzee et al., 2010; Koch et al., 2017; Coetzee et al., 2018). Previous studies have suggested that some species, including *A. novae-zelandiae*, are shared between the two regions (Coetzee et al., 2018). However, results from our community analysis of these same data suggest that no armillarioid species are shared between temperate South America and Australasia (Figure 1) but that morphologically similar species may be present in both geographic regions. Our observation is supported by previous phylogenetic analyses of *Armillaria* species from these regions which shows considerable intraclade substructure from sporocarps in the different biogeographic regions (Pildain et al., 2009) and further suggests that dispersal between the two regions may not be common. This could be due to the closure of overland dispersal routes throughout Antarctica several million years ago, and, as a result, the geographic barrier allowed for two lineages of armillarioid fungi to begin diversification. Our analysis also suggests that Australia and New Zealand are relatively restricted from other geographic regions (Figure 1), following the pattern that dispersal between southern hemisphere biogeographic regions is a much rarer occurrence when compared to the northern hemisphere most likely due to the spatial distance of land masses that have been separated for millions of years. Lastly, we should again point out that our analysis here relies only on the ITS1 and ITS2 markers and to fully assess the level of species distribution and diversity, most notably for the *A. novae-zelandiae* species complex, a thorough multi-gene phylogeny (perhaps including a mating compatibility component) would be needed to assess the level of speciation due to allopatry in the Southern Hemisphere.

Notably, armillarioid species are also depauperate in the lower latitudes—an area characterized by largely tropical climate conditions. For example, *A. puiggarii* is the only known mushroom-forming armillarioid species that appears in tropical South America. Similarly, two to three species were broadly found throughout Africa. Perhaps most interesting are the specimens collected in Southeast Asia, which correspond to one species cluster and show no biogeographic connectedness to other regions (Fig. 1), suggesting that further collecting effort here may uncover a novel suite of species that appear to remain unconnected from other biogeographic regions. A temperate bias in collecting efforts undoubtedly plays a role in this finding as this could certainly be a direct reflection of the disproportionate collecting efforts in temperate regions compared to the tropics. Another explanation could be that species in this lineage are adapted to the cooler and harsher climate in temperate and boreal environments (Koch et al., 2017), or have adapted to temperate woody host species over tropical hosts. A final explanation could be that armillarioid sporocarps are adapted to fruit in cooler environments (Ford et al. 2015), which could explain why they are not observed as frequently in warmer climates. However, it is the tropical rainforests of the Guiana Shield that are the exclusive habitat of the two known sequestrate species of *Guyanagaster* (Henkel et al., 2010; Koch et al., 2017). Given the subhypogeous habit of these sporocarps, it is not implausible that other sequestrate species have yet to be collected and described.

The northern and southern hemispheres have been typically thought to not overlap species distributions (Volk & Burdsall, 1995). While our results are largely consistent with this, there are some exceptions. One of these includes a single *Armillaria* collection from Argentina (species cluster 7 for ITS1 and species cluster 12 for ITS2), which supports a connection from temperate South America with northern hemisphere species. However, whether this species is a relic from a time when certain *Armillaria* species had a much broader distribution, or a recent migrant to the region on imported plant material, remains unknown. When assessed from a 99% clustering threshold, the specimen from Argentina no longer clusters with the northern hemisphere species, suggesting that it does not represent a recent introduction to the region. We also detected several species dominant throughout the northern hemisphere that were also present in South Africa. Previous phylogenetic analyses showed support for at least two introductions of northern hemisphere *Armillaria* species to South Africa which were believed to be introduced centuries ago from potted plants from Europe (Coetzee et al., 2001; Coetzee et al., 2003b; Wingfield et al., 2010). At least one other human-mediated introduction of *Armillaria* from Asia to East Africa has been identified, perhaps introduced to the region on tea plants (Mwenje et al., 2006; Guo et al., 2016). Of the six and five *Armillaria* species clusters at the ITS1 and ITS2 region, respectively, identified from sequences from African specimens, three of them were also represented in samples from the northern hemisphere. This observation supports previous work that there may be three, or more, independent *Armillaria* introductions to Africa. The other two to three *Armillaria* species clusters from Africa include species only present on that continent. This is consistent with our current understanding of the African diversity of *Armillaria*, and if this remains true and no more putatively native *Armillaria* species are discovered, it represents one of the biogeographic zones with the most species-poor representations of *Armillaria*.

### Symbiosis as a driver for distribution and species richness

Another possible contributing factor related to the high species richness in East Asia could be the fact that *Armillaria* species engage in mycorrhizal relationships with mycoheterotrophic plants, most notably the species of achlorophyllus orchids in the genera *Galeola* and *Gastrodia* (Kikuchi & Yamaji, 2010; Guo et al., 2016). Using the dataset of Guo et al. (2016), four of our *Armillaria* species clusters associate with orchids from an ITS1 perspective, and seven species clusters associate with orchids from an ITS2 perspective (Table 1). None of the species clusters are specific to only orchid associates. However, an interesting hypothesis emerges from this association, as to whether this symbiosis is a driver of species diversification, as some mycorrhizal relationships—particularly ectomycorrhizal—are known to enhance diversification of the symbionts (Koide et al., 2008; Wilson et al., 2017; Sato & Toju, 2019). However, to address this question, we need a more thorough understanding of the geographic extent of this interaction across the global mycoheterotrophic orchid distribution. Such research, particularly in tropical regions where these mycoheterotrophic orchids are found, may also lead to the discovery of new *Armillaria* species.

Another symbiosis that may play a role in shaping the distribution of the *Armillaria* species is that with species of the ascomycete genus *Wynnea.* Sclerotia produced by *Wynnea* species, regardless of collecting locality, always contain *Armillaria* hyphae suggesting a dependency of these two lineages on one another (Xu et al., 2019), but it is unclear what is exactly happening in this interaction. Because most of the *Armillaria* sequences from *Wynnea* scloerotia did not pass our filtering thresholds, we independently compared those sequences to our armillarioid species clusters. Armillarioid symbionts of *Wynnea* fell into four species clusters: the clusters 1, 7, 50 and 51 for ITS1 and the clusters 8, 31, 60 and 61 for ITS2. Two of the *Armillaria* symbionts of *Wynnea*, one from West Virginia, USA (species cluster 50 for ITS1 and 60 for ITS2) and the other from Brazil (species cluster 51 for ITS1 and 61 for ITS2), are not conspecific with any of the *Armillaria* species clusters detected in any study and may represent new species. The other two species clusters where *Wynnea* symbionts were detected were those that have large geographic distributions. For species cluster 1, which includes one sporocarp from Central America, there are two additional sequences from *Wynnea* symbionts, collected from Guatemala and Costa Rica. In species cluster 7, the *Wynnea* symbiont from Brazil extends the geographic distribution of this largely East Asian cluster to tropical South America. One possibility for why some *Wynnea* symbionts were collected in locations that are largely disjunct from where the majority of the sporocarps were collected may have to do with the specialization of the sclerotium environment. It is possible that certain armillarioid species were once more widespread or abundant than they currently are but were outcompeted by better adapted armillarioid species. *Wynnea* symbionts may have been buffered from local extinction, particularly in Central and South America, due to the extreme specialization of *Armillaria* species for the *Wynnea* environment. However, an increase in the number of voucher collections of *Wynnea* sclerotia and armillarioid fruiting bodies will be needed to fully test this hypothesis. If this hypothesis is supported, *Wynnea* sclerotia could offer a glimpse into the previous distribution of armillarioid species.

In the *Polyporus umbellatus* interaction, all the *Armillaria* isolates collected by Xing et al. (2017) fell into five and six species clusters when using both ITS1 and ITS2 marker regions, respectively (Table 1). With the ITS1 marker, there is one armillarioid species cluster composed of sequences only known from the association with *P. umbellatus*, suggesting that this may be a source for new *Armillaria* species. This is a similar situation with the putative mycoparasitic interaction between *E. abortivum* and *Armillaria* (Lindner Czederpiltz et al., 2001; Koch & Herr, 2021). In this interaction, the role of symbiosis in driving the distribution and species richness is unclear, often because methods other than the use of ITS region sequencing were used to identify the *Armillaria* associate. Therefore, we do not have a precise understanding of the species that are involved in this interaction. Instead, the distribution of this interaction seems to be driven more so by the distribution of *E. abortivum*, which is restricted to eastern North America and eastern Asia.

Finally, as was previously mentioned, the South American tropics have only a single known mushroom-forming armillarioid species, *A. puiggarii*, but do have two known species of the sequestrate genus *Guyanagaster*. Phylogenetic and biogeographic analyses show that *Guyanagaster* represents the earliest diverging lineage of the armillarioid clade (Koch et al., 2017). This finding was particularly illuminating because sequestration is a unidirectional process that has never been reversed (Hibbett et al., 1997), suggesting that the ancestor of *Guyanagaster* was a gilled mushroom with widespread distribution (Koch et al., 2017). We hypothesized that, because *Guyanagaster* is the only extant lineage, that it is highly specialized for its environment and, as a result, was buffered from extinction unlike its mushroom progenitors (Koch et al., 2017). Further studies have suggested the symbiosis with wood-feeding termites may play a role in its dispersal and ecosystem specialization and, therefore, its persistence (Koch & Aime, 2018; Koch et al., 2021).

## CONCLUSION

In the present study, we used publicly available data to address the global species richness of the genus *Armillaria sensu lato*, a group of fungi most notably recognized as mushroom-forming plant pathogens. By mining public data, we estimate that the overall richness of taxa ranges from 50 to 60 species. The highest species richness was observed in Eastern Asia. We also assessed the overlap of species across geographic regions. We proposed numerous hypotheses regarding the drivers of variability in species diversity and richness between different biogeographic regions and along with symbiotic associations with other fungi and host species. Lastly, we show a proof of concept for the use of publicly available sequence data, from individual voucher specimens and from community sequencing efforts, in understanding fungal species richness in a global context.

## Data and code availability

We used publicly available data as the basis for this study. The public data we used, along with all other associated data that we curated for this study, as well as code for data analysis, can be found at https://github.com/HerrLab/Koch_Global_Armillaria_2021.

## Acknowledgements

We want to thank the anonymous reviewers who provided feedback and comments on the initial submission of this manuscript. This work was completed using the Holland Computing Center of the University of Nebraska, which receives support from the Nebraska Research Initiative. This research was directly supported by start-up funding from the University of Nebraska Agricultural Research Division and the University of Nebraska Office of Research and Economic Development. Additionally, JRH acknowledges funding from the US National Air & Space Administration (Grant #80NSSC17K0737), the US National Science Foundation (EPSCoR Grant #1557417), and US National Institute of Justice (Grant #2017-IJ-CX513-0025), all of which indirectly supported this research. Funding agencies had no role in study design, data collection and interpretation, or the decision to submit the work for publication.

## Author Contributions

RAK and JRH initiated the work, analyzed the data, designed the figures, and drafted the manuscript.

## REFERENCES

Alveshere, B., Bennett, P., Kim, M.S., Klopfenstein, N.B. and LeBoldus, J.M., 2021. First report of *Armillaria cepistipes* causing root disease on *Populus trichocarpa* (black cottonwood) in Oregon, USA. Plant Disease DOI: 10.1094/PDIS-09-20-1993-PDN

Anderson, J. B., and Ullrich, R. C. (1979). Biological species of *Armillaria mellea* in North America. Mycologia 71, 402–414.

Antonín V., Stewart J. E., Ortiz R. M., Kim M.-S., Bonello P. E., Tomšovský M., Klopfenstein N.B. (2021) *Desarmillaria caespitosa*, a North American vicariant of *D. tabescens*, Mycologia 113:4, 776–790 DOI:10.1080/00275514.2021.1890969

Banik, M. T., and Burdsall, H. H. (1998). Assessment of compatibility among *Armillaria cepistipes, A. sinapina*, and North American biological species X and XI, using culture morphology and molecular biology. Mycologia 90, 798–805.

Baumgartner, K., Coetzee, M. P. A., and Hoffmeister, D. (2011). Secrets of the subterranean pathosystem of *Armillaria*. Mol. Plant Pathol. 12, 515–534.

Bengtsson-Palme, J., Ryberg, M., Hartmann, M., Branco, S., Wang, Z., Godhe, A., et al. (2013). Improved software detection and extraction of ITS1 and ITS2 from ribosomal ITS sequences of fungi and other eukaryotes for analysis of environmental sequencing data. Methods Ecol. Evol. 4, 914–919.

Bérubé, J. A., and Dessureault, M. (1988). Morphological characterization of *Armillaria ostoyae* and *Armillaria sinapina* sp. nov. Can. J. Bot. 66, 2027–2034.

Bérubé, J. A., and Dessureault, M. (1989). Morphological studies of the *Armillaria mellea* complex: two new species, *A. gemina* and *A. calvescens*. Mycologia 81, 216–225.

Bruns, T.D., Blackwell, M., Edwards, I., Taylor, A.F., Horton, T., Zhang, N., Kõljalg, U., May, G., Kuyper, T.W., Bever, J.D. and Gilbert, G., 2008. Preserving accuracy in GenBank. Science, 319 (5870).

Cha, J. Y., Sung, J. M., and Igarashi, T. (1994). Biological species and morphological characteristics of *Armillaria mellea* complex in Hokkaido: *A. sinapina* and two new species, *A. jezoensis* and *A. singula*. 1994. Mycoscience 35, 39–47.

Chillali, M., Wipf, D., Guillaumin, J.-J., Mohammed, C., and Botton, B. (1998). Delineation of the European *Armillaria* species based on the sequences of the internal transcribed spacer (ITS) of ribosomal DNA. New Phytol. 138, 553–561.

Coetzee, M. P. A., Wingfield, B. D., Harrington, T. C., Dalevi, D., Coutinho, T. A., and Wingfield, M. J. (2000). Geographical diversity of *Armillaria mellea* s. s. based on phylogenetic analysis. Mycologia 92, 105–113.

Coetzee, M. P. A., Wingfield, B. D., Harrington, T. C., Steimel, J., Coutinho, T. A., and Wingfield, M. J. (2001). The root rot fungus *Armillaria mellea* introduced into South Africa by early Dutch settlers. Mol. Ecol. 10, 387–396.

Coetzee, M. P. A., Wingfield, B. D., Bloomer, P., Ridley, G. S., and Wingfield, M. J. (2003a). Molecular identification and phylogeny of *Armillaria* isolates from South America and Indo-Malaysia. Mycologia 95, 285–293.

Coetzee, M. P. A., Wingfield, B. D., Roux, J., Crous, P. W., Denman, S., and Wingfield, M. J. (2003b). Discovery of two northern hemisphere *Armillaria* species on Proteaceae in South Africa. Plant Pathol. 52, 604–612.

Coetzee, M. P. A., Bloomer, P., Wingfield, M. J., and Wingfield, B. D. (2011). Paleogene radiation of a plant pathogenic mushroom. PLoS One 6, e28545.

Coetzee, M. P. A., Wingfield, B. D., Zhao, J., van Coller, S. J., and Wingfield, M. J. (2015). Phylogenetic relationships among biological species of *Armillaria* from China. Mycoscience 56, 530–541.

Coetzee, M. P. A., Wingfield, B. D., and Wingfield, M. J. (2018). *Armillaria* Root-Rot pathogens: species boundaries and global distribution. Pathogens 7, 83.

Camacho C., Coulouris G., Avagyan V., Ma N., Papadopoulos J., Bealer K., et al. (2009) BLAST+: Architecture and applications. BMC Bioinformatics 10(1):1–9

Darmono, T. W., Burdsall, H. H., and Volk, T. J. (1992). Interfertility among isolates of *Armillaria tabescens* in North America. Sydowia 42, 105–116.

Dunne, C. P., Glen, M., Tommerup, I. C., Shearer, B. L., and Hardy, G. E. S. J. (2002). Sequence variation in the rDNA ITS of Australian *Armillaria* species and intra-specific variation in *A. luteobubalina. Australas*. Plant Pathol. 31, 241–251.

Edgar, R. C. (2010). Search and clustering orders of magnitude faster than BLAST. Bioinformatics 26, 2460–2461.

Federhen, S. (2012). The NCBI Taxonomy database. Nucleic Acids Res. 40, D136–D143.

Ford, K. L., Baumgartner, K., Henricot, B., Bailey, A. M., and Foster, G. D. 2015. A reliable *in vitro* fruiting system for *Armillaria mellea* for evaluation of *Agrobacterium tumefaciens*. Fungal Biol. 119: 859–869.

Garraway, M. O., Hütterman, A., and Wargo, P. M. (1991). “Ontogeny and physiology,” in Armillaria Root Disease Agriculture Handbook 691, ed. C. G. Shaw and G. A. Kile (Washington DC: USDA Forest Service) 21–47.

Guillaumin, J.-J., Mohammed, C., Anselmi, N., Courtecuisse, R., Gregory, S. C., Holdenrieder, O., et al. (1993). Geographical distribution and ecology of the *Armillaria* species in western Europe. Eur. J. Plant Pathol. 23, 321–341.

Guo, S., and Xu, J. (1992). Nutrient source of sclerotia of *Grifola umbellata* and its relationship to *Armillaria mellea*. Acta Bot. Sin. 34, 576–580.

Guo, T., Wang, H. C., Xue, W. C., Zhao, J., and Yang, Z. L. (2016). Phylogenetic analyses of *Armillaria* reveal at least 15 phylogenetic lineages in China, seven of which are associated with cultivated *Gastrodia elata*. PLoS One 11, e0154794.

Haelewaters, D., Dirks, A. C., Kappler, L. A., Mitchell, J. K., Quijada, L., Vandegrift, R., et al. (2018). A preliminary checklist of fungi at Boston Harbor Islands. Northeast. Nat. 25, 45–76.

Hawksworth, D. L., and Lücking, R. (2017). Fungal diversity revisited: 2.2 to 3.8 million species. Microbiol. Spectr. 5, FUNK-0052-2016.

Heinzelmann, R., Dutech, C., Tsykun, T., Labbé, F., Soularue, J.P. and Prospero, S. (2019) Latest advances and future perspectives in *Armillaria* research. Canadian journal of plant pathology 41(1), pp.1–23.

Henkel, T. W., Smith, M. E., and Aime, M. C. (2010). *Guyanagaster*, a new wood-decaying sequestrate fungal genus related to *Armillaria* (Physalacriaceae, Agaricales, Basidiomycota). Am. J. Bot. 97, 1474–1484.

Herr, J.R., Opik, M. and Hibbett, D.S., 2015. Towards the Unification of Sequence-Based Classification and Sequence-Based Identification of Host-Associated Microorganisms. New Phytologist, 205(1), pp. 27–31.

Hibbett, D. S., Pine, E. M., Langer, E., Langer, G., and Donoghue, M. J. (1997). Evolution of gilled mushrooms and puffballs inferred from ribosomal DNA sequences. Proc. Natl. Acad. Sci. U.S.A. 94, 12002–12006.

Hibbett, D., Abarenkov, K., Kõljalg, U., Öpik, M., Chai, B., Cole, J., Wang, Q., Crous, P., Robert, V., Helgason, T. et al., 2016. Sequence-based classification and identification of Fungi. Mycologia, 108(6), pp.1049–1068.

Hood, I. A., and Ramsfield, T. D. (2016). *Armillaria aotearoa* species nova. N. Z. J. For. Sci. 46, 2.

Katoh, K., and Standley, D. M. (2013). MAFFT multiple sequence alignment software version 7: improvements in performance and usability. Mol. Biol. Evol. 30, 772–780.

Kikuchi, G., and Yamaji, H. (2010). Identification of *Armillaria* species associated with *Polyporus umbellatus* using ITS sequences of nuclear ribosomal DNA. Mycoscience 51, 366–372.

Kile, G. A., McDonald, G. I., and Byler, J. W. (1991). Ecology and disease in natural forests,” in Armillaria Root Disease Agriculture Handbook 691, ed. C. G. Shaw and G. A. Kile (Washington DC: USDA Forest Service) 102–121.

Klopfenstein, N. B., Stewart, J. E., Ota, Y., Hanna, J. W., Richardson, B. A., Ross-Davis, A. L., et al. (2017). Insights into the phylogeny of Northern Hemisphere Armillaria: Neighbor-network and Bayesian analyses of translation elongation factor 1-α gene sequences. Mycologia 109, 75–91.

Koch, R. A., and Aime, M. C. (2018). Population structure of *Guyanagaster necrorhizus* supports termite dispersal for this enigmatic fungus. Mol. Ecol. 27, 2667–2679.

Koch, R. A., and Herr, J. R. 2021. Transcriptomics reveals the putative mycoparasitic strategy of the mushroom *Entoloma abortivum* on species of the mushroom *Armillaria*. mSystems 6:5. mSystems00544-21. On bioRxiv as https://doi.org/10.1101/2021.04.30.442184

Koch, R. A., Wilson, A. W., Séné, O., Henkel, T. W., and Aime, M. C. (2017). Resolved phylogeny and biogeography of the root pathogen *Armillaria* and its gasteroid relative, *Guyanagaster*. BMC Evol. Biol. 17, 33.

Koch, R. A., Yoon, G. M., Arayal, U. K., Lail, K., Amirebrahimi, M., LaButti, K., et al. (2021). Symbiotic nitrogen fixation in the reproductive structures of a basidiomycete fungus. Current Biology. https://doi.org/10.1016/j.cub.2021.06.033

Koide, R.T., Sharda, J.N., Herr, J.R. and Malcolm, G.M., 2008. Ectomycorrhizal fungi and the biotrophy–saprotrophy continuum. New Phytologist, 178(2), pp.230–233.

Kozlov, A. M., Darriba, D., Flouri, T., Morel, B., and Stamatakis, A. (2019). RAxML-NG: a fast, scalable and user-friendly tool for maximum likelihood phylogenetic inference. Bioinformatics 35, 4453–4455.

Kuske, C. R., Hesse, C. N., Challacombe, J. F., Cullen, D., Herr, J. R., Mueller, R. C., et al. (2015). Prospects and challenges for fungal metatranscriptomics of complex communities. Fungal Ecol. 14, 133–137.

Leinonen, R., Sugawara, H., Shumway, M., and International Nucleotide Sequence Database Collaboration. (2010). The sequence read archive. Nucleic Acids Res. 39, D19–D21.

Liimatainen, K., Niskanen, T., Dima, B., Kytövuori, I., Ammirati, J. F., and Frøslev, T. G. (2014). The largest type study of Agaricales species to date: bringing identification and nomenclature of Phlegmacium (Cortinarius) into the DNA era. Persoonia 33, 98–140.

Lindner Czederpiltz, D. L., Volk, T. J., and Burdsall, H. H., Jr. (2001). Field observations and inoculation experiments to determine the nature of the carpophoroids associated with *Entoloma abortivum* and *Armillaria*. Mycologia 93, 841–851.

Maphosa, L., Wingfield, B. D., Coetzee, M. P. A., Mwenje, E., and Wingfield, M. J. (2006). Phylogenetic relationships among *Armillaria* species inferred from partial elongation factor 1-alpha DNA sequence data. Australas. Plant Pathol. 35, 513–520.

Mwenje, E., Wingfield, B. D., Coetzee, M. P. A., Nemato, H., and Wingfield, M. J. (2006). *Armillaria* species on tea in Kenya identified using isozyme and DNA sequence comparisons. Plant Pathol. 55, 343–350.

Mysara M., Vandamme P., Props R., Kerckhof F.-M., Leys N., Boon N., Raes J., Monsieurs P. (2017). Reconciliation between operational taxonomic units and species boundaries. FEMS Microbiology Ecology, Volume 93, Issue 4, fix029 doi:10.1093/femsec/fix029

Nilsson, R. H., Larsson, K.-H., Taylor, A. F. S., Bengtsson-Palme, J., Jeppesen, T. S., Schigel, D., et al. (2018). The UNTE database for molecular identification of fungi: handling dark taxa and parallel taxonomic classifications. Nucleic Acids Res. 47, D259–D264.

Oliveira, J. J. S., Moncalvo, J.-M., Margaritescu, S., and Capelari, M. (2020). A morphological and phylogenetic evaluation of *Marasmius* sect. *Globulares* (Globulares-Sicci complex) with nine new taxa from the Neotropical Atlantic Forest. Persoonia 44, 240–277.

Ota, Y., Matsushita, N., Nagasawa, E., Terashita, T., Fukuda, K., and Suzuki, K. (1998). Biological species of *Armillaria* in Japan. Plant Dis. 82, 537–543.

Pildain, M. B., Coetzee, M. P. A., Rajchenberg, M., Petersen, R. H., Wingfield, M. J., and Wingfield, B. D. (2009). Molecular phylogeny of *Armillaria* from the Patagonian Andes. Mycol. Prog. 8, 181–194.

Qin, G. F., Zhao, J., and Korhonen, K. (2007). A study on intersterility groups of *Armillaria* in China. Mycologia 99, 430–441.

R Core Team (2013). R: A language and environment for statistical computing. R Foundation for Statistical Computing, Vienna, Austria. http://www.R-project.org/.

Rognes, T., Flouri, T., Nichols, B., Quince, C., and Mahé, F. (2016). VSEARCH: a versatile open source tool for metagenomics. PeerJ 4, e2584.

Sato, H., and Toju, H. (2019). Timing of evolutionary innovation: scenarios of evolutionary diversification in a species-rich fungal clade, Boletales. New Phytol. 222, 1924–1935.

Schoch, C. L., Seifert, K. A., Huhndorf, S., Robert, V., Spouge, J. L., Levesque, C. A., Chen, W., et al. 2012. Nuclear ribosomal internal transcribed spacer (ITS) region as a universal DNA barcode marker for *Fungi*. Proc. Natl. Acad. Sci. U.S.A. 109, 6241–6246.

Sipos, G., Prasanna, A.N., Walter, M.C., O’Connor, E., Bálint, B., Krizsán, K., Kiss, B., Hess, J., Varga, T., Slot, J. and Riley, R., 2017. Genome expansion and lineage-specific genetic innovations in the forest pathogenic fungi *Armillaria*. Nature ecology & evolution, 1(12), pp.1931–1941.

Smith, M. L., Bruhn, J. N., and Anderson, J. B. (1992). The fungus *Armillaria bulbosa* is among the largest and oldest living organisms. Nature 356, 428–431.

Tedersoo, L., Bahram, M., Põlme, S., Kõljalg, U., Yorou, N. S., Wijesundera, R., et al. (2014). Global diversity and geography of soil fungi. Science 346, 1256688.

Truong, C., Mujic, A. B., Healy, R., Kuhar, F., Furci, G., Torres, D., et al. (2017). How to know the fungi: combining field inventories and DNA-barcoding to document fungal diversity. New Phytol. 214, 913–919.

Tsykun, T., Rigling, D., and Prospero, S. (2013). A new multilocus approach for a reliable DNA-based identification of *Armillaria* species. Mycologia 105, 1059–1076.

Volk, T. J., and Burdsall, H. H., Jr. (1995). A Nomenclatural Study of Armillaria and Armillariella Species. Oslo: Synopsis Fungorum.

Vu, D., Groenewald, M., de Vries, M., Gehrmann, T., Stielow, B., Eberhardt, U., et al. (2019). Large-scale generation and analysis of filamentous fungal DNA barcodes boosts coverage for kingdom fungi and reveals thresholds for fungal species and higher taxon delimation. Stud. Mycol. 92, 135–154.

Watling, R., Kile, G. A., and Burdsall, H. H., Jr. (1991). “Nomenclature, taxonomy, and identification,” in Armillaria Root Disease Agriculture Handbook 691, ed. C. G. Shaw and G. A. Kile (Washington DC: USDA Forest Service) 1–9.

Wilson, A. W., Hosaka, K., and Mueller, G. M. (2017). Evolution of ectomycorrhizas as a driver of diversification and biogeographic patterns in the model mycorrhizal mushroom genus *Laccaria*. New Phytol. 213, 1862–1873.

Wingfield, M. J., Coetzee, M. P. A., Crous, P. W., Six, D., and Wingfield, B. D. (2010). Fungal phoenix rising from the ashes? IMA Fungus 1, 149–153.

Wright, E. S. (2016). Using DECIPHER v2.0 to analyze big biological sequence data in R. R J. 8, 352–359.

Xing, X., Men, J., and Guo, S. (2017). Phylogenetic constrains on *Polyporus umbellatus-Armillaria* associations. Sci. Rep. 7, 4226.

Xu, F., LoBuglio, K. F., and Pfister, D. H. (2019). On the co-occurrence of species of *Wynnea (Ascomycota, Pezizales, Sarcoscyphaceae)* and *Armillaria (Basidiomycota, Agaricales, Physalacriaceae)*. Fungal Syst. Evol. 4, 1–12.

Yu, G., Tsan-Yuk Lam, T., Zhu, H., and Guan, Y. 2018. Two methods for mapping and visualizing associated data on phylogeny using Ggtree. Mol. Biol. Evol. 35, 3041–3043.

Zhao, J., Dai, Y. C., Qin, G. F., Wei, Y. L., Wang, H. C., and Yuan, H. S. (2008). New biological species of *Armillaria* from China. Mycosystema 27, 156–170.

Zolciak, A., Bouteville, R.-J., Tourvieille, J., Roeckel-Drevet, P., Nicolas, P., and Guillaumin, J.-J. (1997). Occurrence of *Armillaria ectypa* (Fr.) Lamoure in peat bogs of the Auvergne—the reproduction system of the species. Cryptogam. Mycol. 18, 299–313.

